# Sex hormones gate the neuronal control of dendritic cell migration and immunity

**DOI:** 10.64898/2026.10.01.754550

**Authors:** Konrad Knöpper, Tamar L Ben-Shaanan, Jinping An, Ying Xu, Jason G. Cyster

**Author notes:** **Correspondence:** (K.K.), (J.G.C.).

## Abstract

How endocrine states modulate local immune responses is a fundamental question in immunology. Sexual dimorphism is a striking example: females generally mount stronger immune responses than males, yet how sex hormones intersect with local signals is poorly understood. Here, we identify an endocrine-neuro–immune circuit in which sex hormones determine whether immune cells respond to nociceptor signals. In female skin, nociceptor-derived CGRP promotes dendritic cell (DC) migration and T cell priming indirectly through inflammatory monocytes. CGRP induces monocyte production of Activin A, which upregulates CXCR4 on DCs and promotes migration to draining lymph nodes. In males, androgen receptor signaling suppresses monocyte Activin A and silences this circuit. Disrupting the relay abolishes enhanced female immunity, whereas deleting androgen receptor in male monocytes restores neuronal control. Thus, sex hormones regulate immunity by determining how immune cells interpret neuronal signals, revealing how endocrine state can gate local neuroimmune communication to generate sex-specific immune responses.

**Highlights:**

- Nociceptor neurons enhance dendritic cell migration selectively in females
- Neuronal CGRP signals through monocytes to promote dendritic cell migration
- Monocyte-derived Activin A induces CXCR4 on dendritic cells
- Androgen signaling gates neuronal control of immunity in males

**In Brief:** Knöpper et al. show that nociceptor neurons drive dendritic cell migration and adaptive immunity in female but not male skin. Neuronal CGRP acts through recruited monocytes, which produce Activin A to induce CXCR4 on dendritic cells. Androgen signaling in monocytes silences this circuit in males.

## Introduction

Barrier tissues continuously integrate environmental information with signals from resident epithelial, immune, and neuronal cells to mount protective responses while limiting immunopathology^1–6^. In the skin, dendritic cells (DCs) occupy a central position in this process. Following antigen encounter and inflammatory activation, dermal DCs migrate to skin-draining lymph nodes, where they initiate and shape adaptive immune responses^7,8^. DCs are instructed by local inflammatory cues and tissue-derived signals to promote appropriate T helper cell differentiation and effector responses^9^. Dysregulation of these pathways underlies skin diseases, including contact dermatitis, a common allergic reaction affecting ∼20% of the general population^10^, which is characterized by exaggerated type 2 immune responses^11^. Small reactive chemicals called haptens cause contact dermatitis in humans, and are well established to study this condition in mice^12^.

A striking feature of immune responses across tissues is their sexual dimorphism^13,14^. Females generally mount stronger innate and adaptive immune responses than males, contributing to enhanced protection against infection and vaccination efficacy but also increased susceptibility to autoimmune and allergic diseases^15–17^. In the skin, sex differences are particularly pronounced, with females displaying heightened immune activation, increased DC numbers and enhanced T cell responses following antigen challenge^18^. Contact dermatitis shows female biased prevalence in mutilple studies^19,20^. Allthough allergen exposure patterns have been associated with a female bias in developing contact dermatitis^21^, controlled exposure tests provide evidence for an intrinsic sex difference^22^. Sex hormones and sex chromosome-linked genes can regulate immune-cell development and function, yet how these systemic determinants interact with signals generated locally within tissues remains poorly understood. In particular, whether sex influences how immune cells interpret tissue-derived signals, rather than simply altering their abundance or basal state, is unclear.

The skin is densely innervated by sensory neurons that detect environmental and inflammatory stimuli and communicate rapidly with surrounding cells^23–25^. Sensory neurons, including those expressing the transient receptor potential vanilloid 1 (TrpV1) channel, also classified as nociceptors, can directly sense noxious stimuli and allergens, leading to the release of neuropeptides such as calcitonin gene–related peptide (CGRP) and substance P ^26^. These neuro-immune interactions have emerged as key regulators of cutaneous immunity, influencing DC activation, migration and T cell priming^27–32^. Notably, sensory neurons can act as primary sensors of allergenic stimuli, relaying signals that shape downstream immune responses^27,33^. However, whether neuronal control of immunity is itself subject to sexual dimorphism, and how such differences might contribute to sex-specific immune outcomes in the skin, has not been studied so far.

Here, we identify an endocrine-neuro–immune circuit that controls sexually dimorphic skin immunity. We show that nociceptor neurons promote DC migration and adaptive immune activation in a sex-dependent manner and uncover a myeloid relay through which neuronal signals are transmitted to DCs. We further establish that androgen signaling acts within this relay to determine responsiveness to neuronal input. These findings reveal a mechanism by which endocrine state gates neuroimmune communication, allowing the same tissue-derived neuronal signal to produce distinct immune outcomes in males and females.

## Results

### Female skin mounts stronger adaptive immune responses

Given the known sexual-dimorphism in skin immunity and open questions about molecular mechanisms, we investigated male-female differences in contact hypersensitivity (CHS) models. Consistent with previous observations^14,18,22^, we found that female mice mount a stronger cutaneous immune response than males. Using the FITC hapten CHS model, topical application of FITC-dibutylphthalate induced significantly greater tissue swelling during the challenge phase in females (Fig. 1A, B), accompanied by increased accumulation of CD4⁺ T cells in the skin (Fig. 1C). To test whether the enhanced response originated during the T cell priming phase, we immunized male and female mice with several immune stimuli, namely FITC-dibutylphthalate, heat-killed *Listeria monocytogenes* (HKLM), and aluminum hydroxide, along ovalbumin (Ova) as an antigen in each case. We measured the primary CD4 T cell response by quantifying transferred Ova-specific OT-II TCR transgenic T cells in the skin draining lymph node (sdLN). FITC and HKLM responses were analyzed 2 days post treatment to quantify OT-II T cell proliferation, while alum treated mice were analyzed at day 5 to measure cytokine profiles in addition to OT-II T cell expansion. Across all stimuli, females consistently exhibited increased OT-II T cell proliferation and accumulation relative to males (Fig. 1D-F). The cytokine profile of the T cells was not altered comparing sexes (Fig. S1A). Thus, enhanced female skin immunity is already evident during the priming phase and is conserved across distinct inflammatory stimuli.

**Fig. 1.**
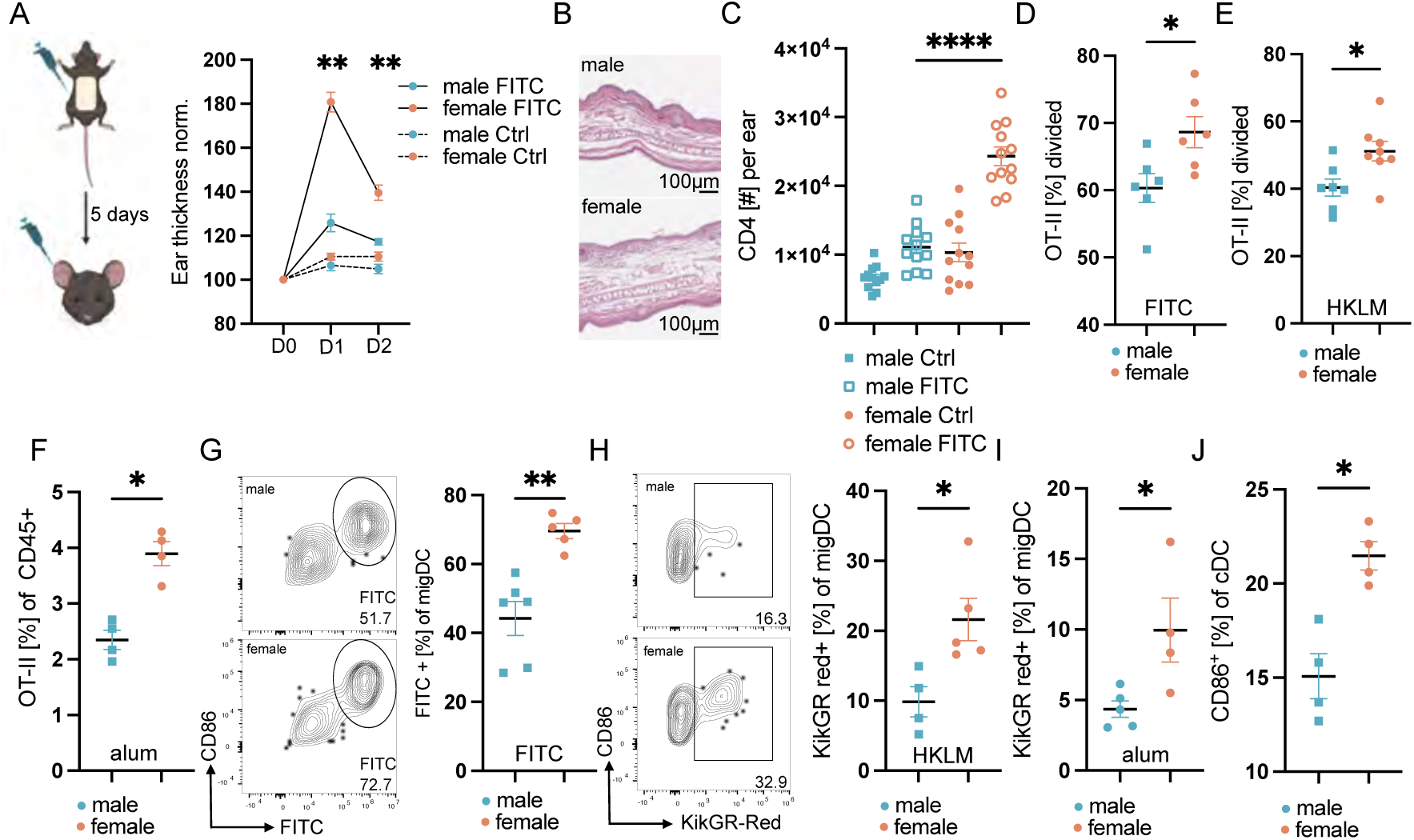
Female skin mounts stronger adaptive immune responses. A) Schematic of experimental set-up to induce CHS response (left). Ear thickness measurement at D0, D1, and D2 post challenge phase, normalized to D0 thickness (right). 6 mice per group are displayed. Figure created with BioRender.com B) Representative H&E histology of ear sections collected at D2 post challenge. C) Quantification of CD4 T cell numbers per ear at D2 post challenge either with hapten (FITC) or solvent alone (Ctrl). D) Quantification of OT-II division based on CTV dilution at D2 post FITC and Ova treatment. E) Quantification of OT-II division based on CTV dilution at D2 post HKLM and Ova treatment. F) Quantification of OT-II accumulation based on frequency of CD45+ cells at D5 post alum and Ova treatment. G) Representative flow cytometry plot of migratory DCs from the sdLN 18 h post FITC treatment (left). Quantification of FITC+ migratory DCs from the sdLN 18 h post FITC treatment (right). H) Representative flow cytometry plot of migratory DCs from the sdLN 18 h post HKLM treatment (left). Quantification of KikGR-Red+ migratory DCs from the sdLN 18h post HKLM treatment (right). I) Quantification of KikGR-Red+ migratory DCs from the sdLN 18 h post alum treatment. J) CD86 expression on skin DC isolated 18 h post FITC treatment of WT mice. All data were pooled from two independent experiments or display one representative experiment of N= 2-6 mice per group and represented as mean ± SEM. \*\*\*\**P*<0.0001, ** *P*<0.01, * *P*<0.05, two-way ANOVA or t-test.

Previously, increased T cell responses in females relative to males was connected to higher skin DC numbers, and an associated increase in DCs reaching the sdLN^18^. Consistent with this, we detected higher numbers of DCs in female skin (Fig. S1B). Tracing the DC migration either via FITC-uptake or KikGR fluorescent protein photo-conversion, we could observe higher frequencies of migrating DCs in the sdLN in females across stimuli (Fig. 1G-I). KikGR photo-conversion efficiency in the skin was confirmed to be equivalent in males and females (Fig. S1C). To test whether DC abundance in the skin was the main driver of increased DC migration in females, we measured DC activation in the skin after stimulation. Surprisingly, we observed already in the skin an increase in activated CD86 expressing DCs comparing females to males (Fig. 1J), hinting towards additional factors imprinting the sex-differences. We therefore asked whether upstream tissue-derived cues could differentially regulate DC behavior between sexes, independent of DC numbers.

### Nociceptor neurons drive female-biased DC migration and T cell priming

The skin is densely innervated by nociceptor neurons that rapidly respond to environmental stimuli and have been shown to directly influence DC activation and migration^27,29,34–36^. Importantly, FITC-dibutylphthalate^31^, bacterial challenge^37^ and aluminum hydroxide^38^ have all been described to activate nociceptor neurons directly or indirectly, among several other stimuli. Given the well-established sexual dimorphism in sensory neuron function^34–36^, we hypothesized that neuronal signaling might represent a previously unrecognized regulator of sex-specific DC migration. To test this possibility, we generated mice lacking nociceptors, which are a subset of sensory neurons crucial for generating pain sensation and the main cutaneous sources for the immune modulating neuropeptides CGRP and substance P^39^, by crossing TrpV1-Cre mice to mice harboring a conditional allele of diphtheria toxin (TrpV1^DTA^). We validated the functionality of the nociceptor neuron deletion via hot-plate assay (Fig. S2A). Importantly, measuring CD86 level in TrpV1^DTA^ mice showed no sex-difference after FITC-dibutylphthalate stimulation (Fig. 2A). Analyzing DC migration via FITC-uptake, as well as photo-conversion revealed that lack of nociceptor neurons reduced female DC migration significantly, while having no obvious effect in males, across the tested stimuli (Fig. 2B-D). Furthermore, the observed male-female differences in WT mice were not present in TrpV1^DTA^ mice (Fig. 2B-D). In addition to tracing migration of DC from the back skin to the sdLN, we also observed a reduction of DC migration from the ear skin to the auricular LN, highlighting that nociceptor neurons promote DC migration at different skin parts (unpubl. obs.). The importance of direct neuronal activation was tested via local blocking of cell depolarization via Qx314^40^. This treatment selectively reduced DC migration in female mice (Fig. 2E). Analyzing the T cell response, we observed reduced responses in female but not male TrpV1^DTA^ mice in accord with the effect on DC migration (Fig. 2F-H). Importantly, DC numbers in the skin were not altered by the absence of nociceptor neurons (Fig. S2B), and their subset distribution^41^ remained intact (Fig. S2C, D), indicating that neuronal input regulates DC function rather than DC abundance. Together, these findings identify nociceptors as critical drivers of the enhanced DC migration and adaptive immune responses observed in females.

**Fig. 2.**
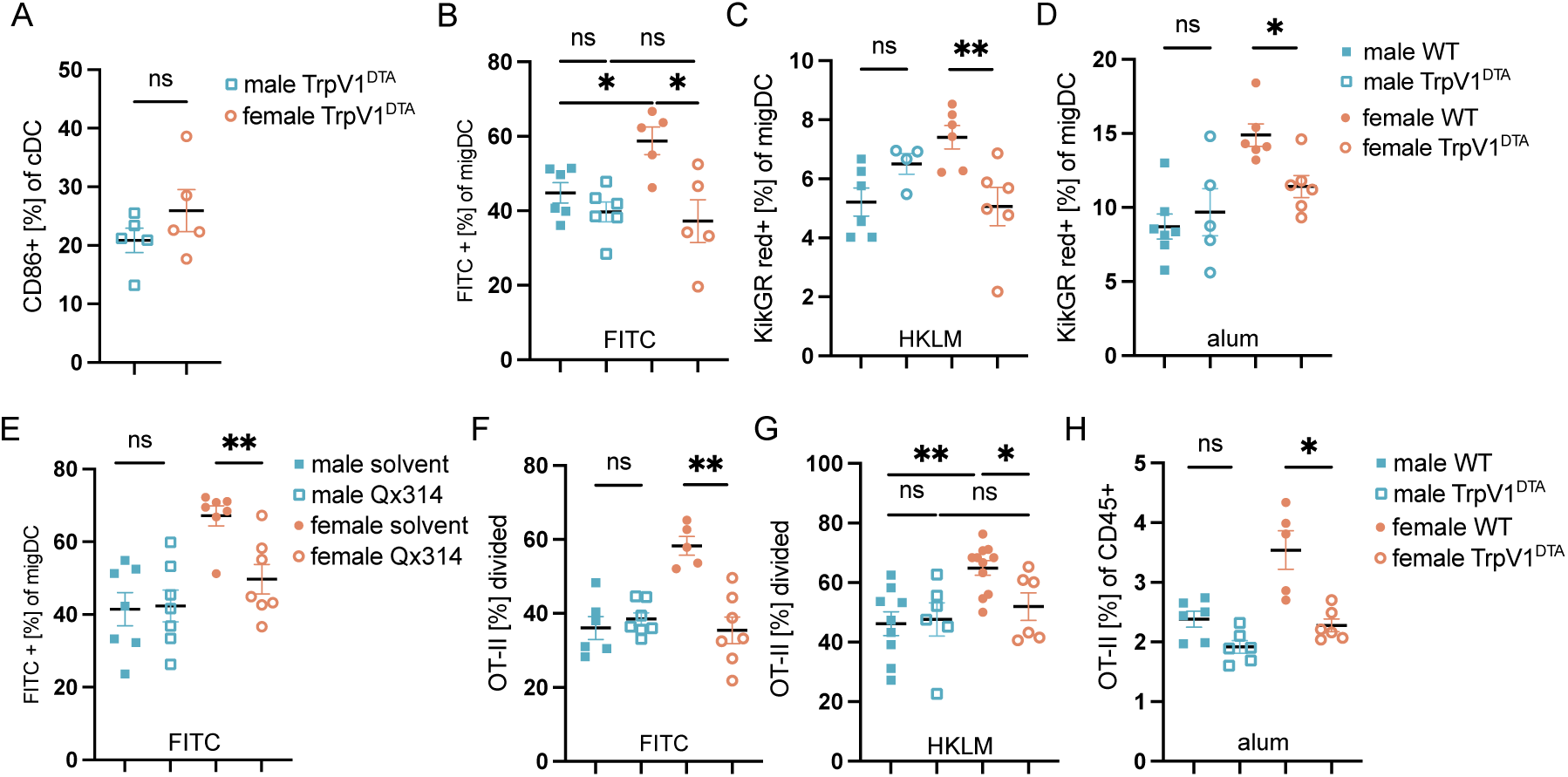
Nociceptor neurons drive female-biased DC migration and T cell priming. A) Quantification of CD86 expression on skin DC isolated 18 h post FITC treatment of TrpV1^DTA^ mice. B) Quantification of FITC+ migratory DCs from the sdLN 18 h post FITC treatment in WT and TrpV1^DTA^ mice. C) Quantification of KikGR-Red+ migratory DCs from the sdLN 18 h post HKLM treatment in WT and TrpV1^DTA^ mice. D) Quantification of KikGR-Red+ migratory DCs from the sdLN 18 h post alum treatment in WT and TrpV1^DTA^ mice. E) Quantification of FITC+ migratory DCs from the sdLN 18 h post FITC. Mice were treated with Qx314 neuronal activation inhibitor or solvent control. F) Quantification of OT-II division based on CTV dilution at D2 post FITC and Ova treatment in WT and TrpV1^DTA^ mice. G) Quantification of OT-II division based on CTV dilution at D2 post HKLM and Ova treatment in WT and TrpV1^DTA^ mice. H) Quantification of OT-II accumulation based on frequency of CD45+ cells at D5 post alum and Ova treatment in WT and TrpV1^DTA^ mice. All data were pooled from two independent experiments or display one representative experiment of N= 2-6 mice per group and represented as mean ± SEM. ** *P*<0.01, * *P*<0.05, ns = non-significant, two-way ANOVA or t-test.

### CGRP is the neuronal mediator of female DC migration

We next sought to define the molecular mechanism by which nociceptors promote DC migration in a sex-dependent manner. Nociceptor neurons rapidly communicate with surrounding cells through the release of neuropeptides, several of which have been implicated in regulating DC behavior^27–29^ (Fig. 3A). We therefore asked whether specific neuronal mediators could account for the enhanced DC migration in females. Substance P has been reported to promote DC migration during papain-induced inflammation^27^, which we could reproduce by blocking Substance P signaling with the antagonist QWF in papain exposed mice (Fig. S3A). However, QWF treatment did not impact DC migration post FITC application (Fig. S3B). Furthermore, we analyzed Tac1 (Substance P) deficient mice and observed no impact on DC migration or T cell response in our models (Fig. S3C, D).

**Fig. 3.**
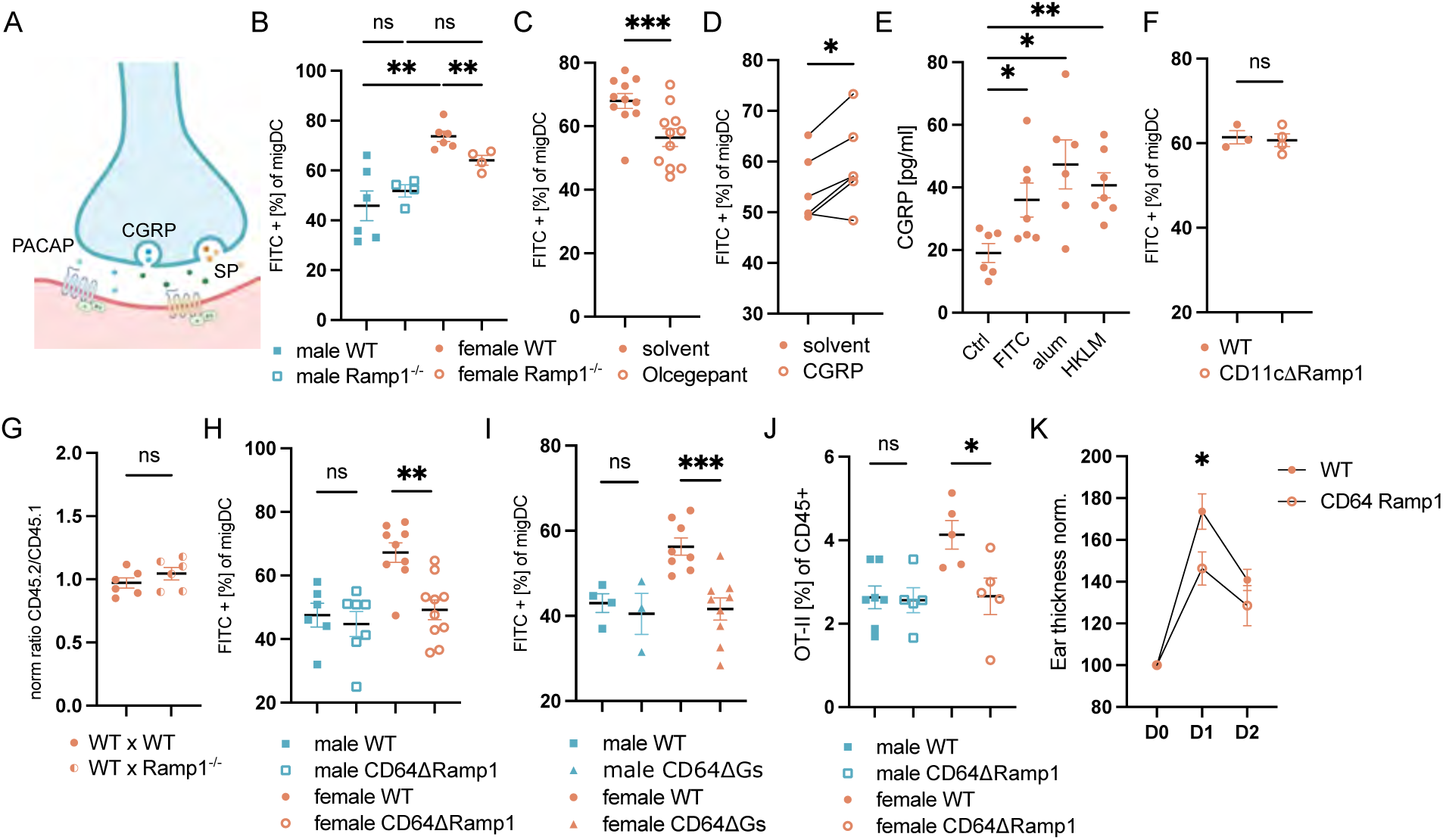
CGRP is the neuronal mediator of female DC migration. A) Schematic representation of neurotransmitter release and signaling on target cell. SP, substance P; PACAP, pituitary adenylate cyclase-activating polypeptide. Figure created with BioRender.com B) Quantification of FITC+ migratory DCs from the sdLN 18 h post FITC treatment in WT and Ramp1^-/-^ mice. C) Quantification of FITC+ migratory DCs from the sdLN 18 h post FITC treatment in female WT mice treated with Olcegepant Ramp1 antagonist or solvent control. D) Quantification of FITC+ migratory DCs from the sdLN 18 h post FITC treatment in TrpV1^DTA^ female mice with or without supplementation of CGRP. E) Quantification of CGRP secretion via ELISA from female skin explants after 30 min of the indicated stimuli. F) Quantification of FITC+ migratory DCs from the sdLN 18 h post FITC treatment in female WT and CD11cΔRamp1 mice. G) Quantification of FITC+ migratory DCs from the sdLN 18 h post FITC treatment in mixed BM chimeric female mice. H) Quantification of FITC+ migratory DCs from the sdLN 18 h post FITC treatment in WT and CD64ΔRamp1 mice. I) Quantification of FITC+ migratory DCs from the sdLN 18 h post FITC treatment in WT and CD64ΔGs mice. J) Quantification of OT-II accumulation based on frequency of CD45+ cells at D5 post alum and Ova treatment in WT and CD64ΔRamp1 mice. K) Ear thickness measurement at D0, D1, and D2 post challenge phase in female WT and CD64ΔRamp1 mice, normalized to D0 thickness. 6 mice per group are displayed. All data were pooled from two to three independent experiments or display one representative experiment of N= 2-5 mice / group and represented as mean ± SEM. *** *P*<0.001, ** *P*<0.01, * *P*<0.05, ns = non-significant, two-way ANOVA or t-test.

We next focused on CGRP, a prominent neuropeptide released by TrpV1⁺ nociceptors. To test its role, we analyzed mice deficient for the CGRP receptor component Ramp1. Strikingly, Ramp1 deficiency selectively reduced DC migration in females (Fig. 3B), partially phenocopying the effect of nociceptor neuron ablation. Local pharmacological inhibition of CGRP signaling using Olcegepant similarly impaired DC migration in females (Fig. 3C), whereas exogenous CGRP administration restored DC migration in TrpV1^DTA^ female mice (Fig. 3D). Importantly, all previously used immune stimuli were able to induce CGRP secretion from skin explants (Fig. 3E). These findings identify CGRP as a key neuronal mediator driving female-biased DC migration.

### CGRP acts indirectly, not on dendritic cells

We next asked whether CGRP acts directly on DCs in our model, since DCs were reported to respond to this neuropeptide in some settings^28,29^. To this end, we crossed Ramp1 floxed and CD11c-Cre mice, generating a DC-specific Ramp1-deficiency (CD11cΔRamp1). Surprisingly, we did not observe a defect in DC migration in CD11cΔRamp1 mice compared to WT, pointing towards a non-DC intrinsic effect of CGRP (Fig. 3F). To substantiate our observation, we generated mixed bone marrow (BM) chimeras using CD45 congenically distinct WT and Ramp1-deficient BM. Analysis of these mice revealed no intrinsic migration defect in Ramp1-deficient DCs (Fig. 3G), suggesting that CGRP signals through an intermediary cell population. To validate irradiation-generated chimeras as a suitable approach, we reconstituted irradiated TrpV1^DTA^ mice with WT BM, which maintained the migration defect in TrpV1^DTA^ mice (Fig. S3E), highlighting that irradiation does not interfere with the neuronal induced DC migration. Together, the data indicate that CGRP does not act directly on DCs to promote their migration.

### CD64⁺ monocytes are the cellular relay for CGRP signaling

To identify the relevant target cell population, we next deleted Ramp1 in CD64⁺ monocytes and macrophages (CD64ΔRamp1), cells that have been reported to be CGRP responsive^3,42,43^. This resulted in a marked reduction in DC migration selectively in females (Fig. 3H). Furthermore, deletion of Gs, a key signaling protein downstream of the CGRP receptor, in the same cells (CD64ΔGs) impaired DC migration in females (Fig. 3I), demonstrating that CGRP signaling in CD64^+^ cells is required for this process. As an independent genetic test, we deleted Ramp1 using Ms4a3-Cre, a second Cre driver targeting monocyte/macrophage-lineage cells. Ms4a3-mediated Ramp1 deletion similarly reduced DC migration in females (Fig. S3F). Together with the absence of a migration defect following DC-specific Ramp1 deletion (Fig. 3F), these findings support an indirect mechanism in which CGRP acts through monocytes/macrophages rather than directly on DCs. Notably, cell type-restricted deletion of Ramp1 or Gs produced a more pronounced reduction in DC migration than global Ramp1 deficiency. This difference suggests that CGRP signaling in additional cell populations may exert opposing effects on DC migration.

Functionally, loss of Ramp1 in CD64⁺ cells reduced antigen-specific T cell expansion post aluminum hydroxide immunization (Fig. 3J), as well as ear swelling in the CHS model (Fig. 3K), establishing that CGRP-dependent signaling in CD64^+^ cells drives both DC migration and downstream adaptive immunity. Together, these findings indicate that neuronal control of DC migration is mediated indirectly through CGRP signaling in CD64⁺ myeloid cells.

We next sought to define how CGRP signaling in CD64⁺ myeloid cells promotes DC migration. To this end, we performed single-cell RNA sequencing of CD64⁺ cells isolated from the skin of WT and CD64ΔRamp1 female mice under steady-state condition or 8 hours after FITC treatment. This analysis revealed marked cellular heterogeneity within the CD64⁺ compartment (Fig. 4A, Fig. S4A–B) and a substantial shift in population composition upon activation (Fig. 4B). Most prominently, we observed an influx of Ly6C^high^ monocytes expressing low levels of CD64 into the skin following stimulation (Fig. 4B), which we confirmed by flow cytometry (Fig. 4C). We did not observe a difference in monocyte recruitment comparing WT with Ramp1-deficient mice at this early time point (Fig. 4B, C) in contrast to previously observed increased accumulation of Ramp1-deficient monocytes in inflamed skin at later time points^3^.

**Fig. 4.**
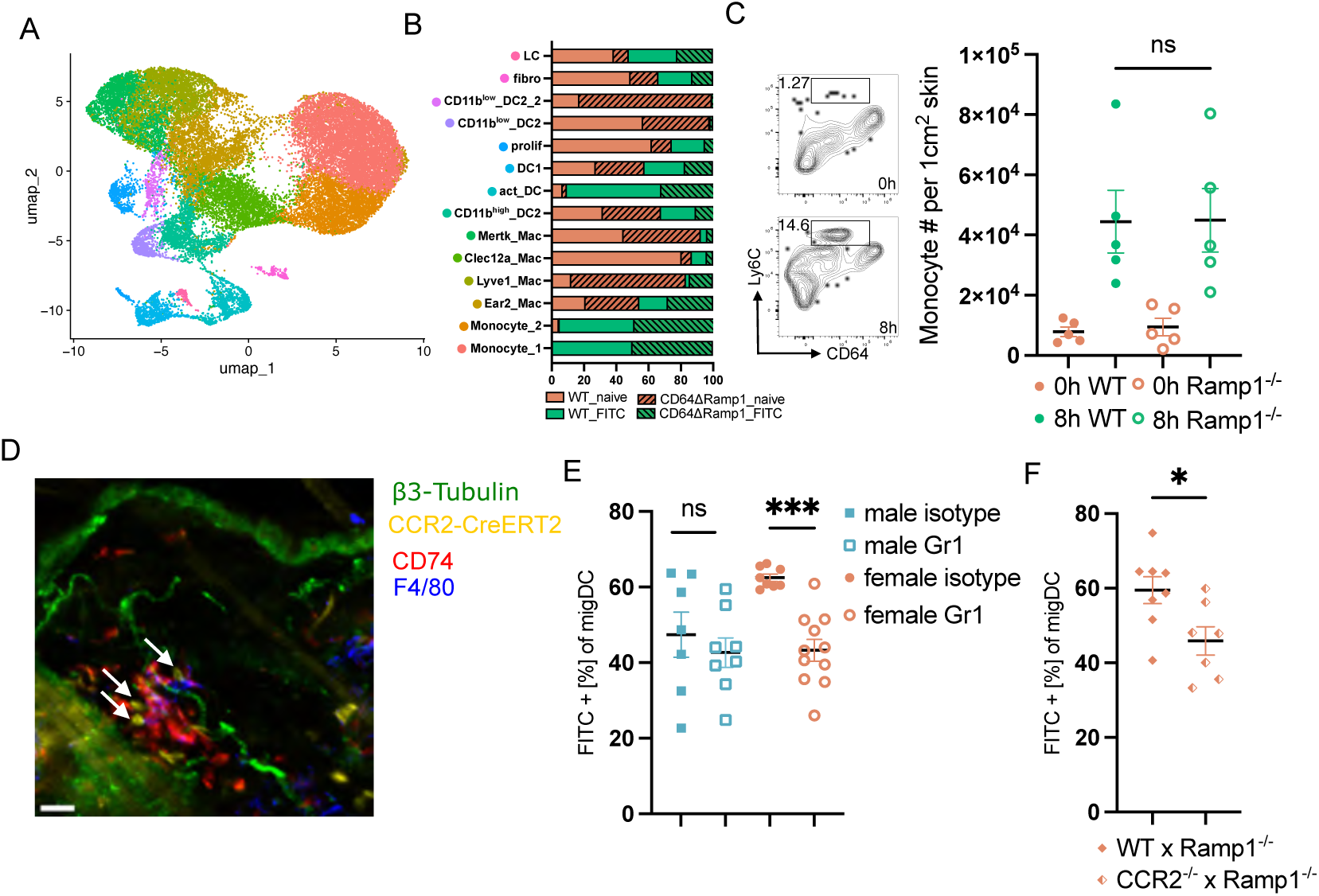
CD64⁺ monocytes are the cellular relay for CGRP signaling. A) UMAP projection displaying the clustering of sorted CD64+ immune cells based on scRNA-seq data, color represents cluster identity. B) Relative cluster representation across analyzed conditions, normalized to cell number per condition. C) Representative flow cytometry plot (left) and quantification (right) of Ly6C^high^ monocyte infiltration into the skin of female mice 8 h post FITC application. D) Representative IF image of skin 8 h post dibutylphthalate application. βIII-tubulin (green), CCR2CreERT2-Ai14 (yellow), CD74 (red), F4/80 (blue). Scale bar: 20 µm. E) Quantification of FITC+ migratory DCs from the sdLN 18 h post FITC treatment in WT pre-treated either with isotype control or Gr1-depletion antibody. F) Quantification of FITC+ migratory DCs from the sdLN 18 h post FITC treatment in mixed BM chimeric female mice. All data were pooled from two to three independent experiments or display one representative experiment of N= 3-5 mice / group and represented as mean ± SEM. *** *P*<0.001, * *P*<0.05, ns = non-significant, two-way ANOVA or t-test.

Since recruited monocytes were found in close proximity to DCs and neurons in the skin (Fig. 4D), we asked whether these Ly6C^+^ cells mediate the CGRP-dependent enhancement of DC migration. Depletion of Ly6C⁺ cells using Gr1 antibody resulted in a significant reduction in DC migration to sdLNs in females (Fig. 4E). Because Gr1 targets both monocytes and neutrophils (via Ly6C and Ly6G, respectively), we next selectively depleted neutrophils using anti-Ly6G antibody, which had no effect on DC migration (Fig. S4C), indicating a specific role for monocytes.

To further test the role of Ramp1 in monocytes, we generated mixed Ramp1^-/-^:CCR2^-/-^ BM chimeras in which CCR2-deficiency prevented recruitment of Ramp1 proficient monocytes to inflamed skin (Fig. S4D), practically generating a monocyte specific Ramp1-deficiency in the inflamed skin. Under these conditions, DC migration was significantly reduced in females (Fig. 4F), confirming that CGRP signaling in recruited monocytes is required for this process.

### Monocyte-derived Activin A induces CXCR4 to promote DC migration

We next sought to identify monocyte-derived factors that promote DC migration downstream of CGRP signaling. Differential gene expression analysis highlighted *Inhba* as a candidate mediator, as it was selectively expressed in monocytes (Fig. 5A) and enriched in WT compared to CD64ΔRamp1 cells (Fig. 5B), while its receptor subunits *Acvr2a* and *Acvr1b* were expressed on DC (Fig. S5A, B)^44^. Public transcriptomic dataset analysis showed that *Inhba* expression is inducible by CGRP or dorsal root ganglia (DRG) derived factors in myeloid cells (Fig. S5C, D)^43,45^.

**Fig. 5.**
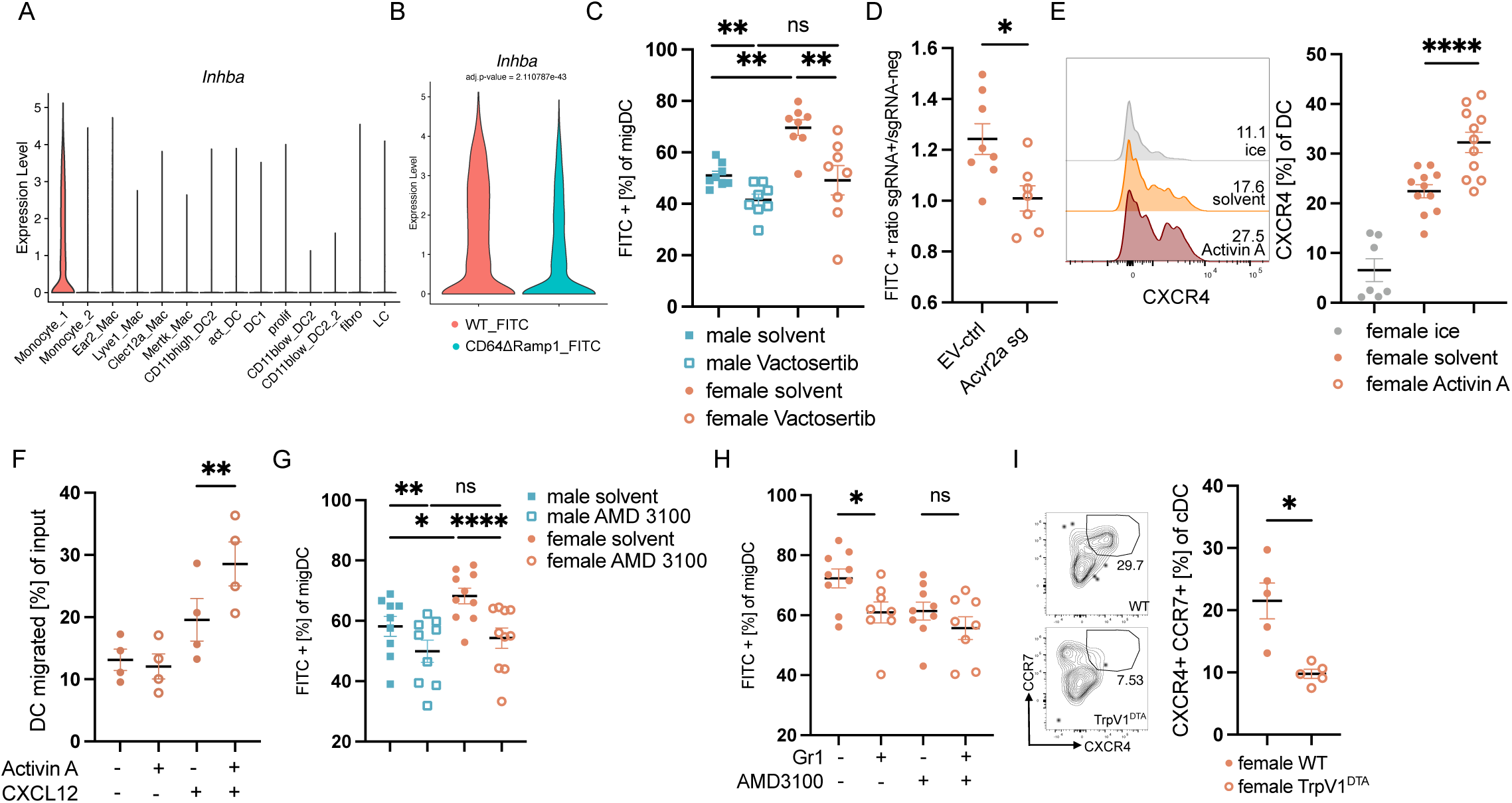
Monocyte-derived Activin A induces CXCR4 to promote DC migration. A) Quantification of *Inhba* normalized expression across clusters from scRNA-seq data. B) Normalized expression of *Inhba* gene in cluster Monocyte_1 comparing WT and CD64ΔRamp1 mice. C) Quantification of FITC+ migratory DCs from the sdLN 18 h post FITC treatment in WT treated either with solvent control or Vactosertib Activin A receptor antagonist. D) Quantification of FITC+ migratory DCs from the sdLN 18 h post FITC treatment in mice transduced with *Acvr2a* targeting sgRNA or empty vector (EV) control. E) Representative flow cytometry histogram displaying CXCR4+ percentage (left) and quantification (right) of CXCR4 expression on female skin DCs after *ex vivo* treatment with Activin A for 4 h. F) Quantification of female splenic DC migration towards CXCL12 after 4 h pre-incubation with Activin A. G) Quantification of FITC+ migratory DCs from the sdLN of male and female mice 18 h post FITC treatment in WT treated either with solvent or AMD3100 CXCR4 antagonist. H) Quantification of FITC+ migratory DCs from the sdLN 18 h post FITC treatment in female WT treated either with Gr1-depletion antibody and AMD3100. I) Representative flow cytometry plot (left) and quantification (right) of CXCR4 and CCR7 positive female skin DCs 18 h post FITC application. All data were pooled from two to three independent experiments or display one representative experiment of N= 3-5 mice / group and represented as mean ± SEM. \*\*\*\**P*<0.0001, ** *P*<0.01, * *P*<0.05, ns = non-significant, two-way ANOVA or t-test.

*Inhba* encodes Activin A, a homodimeric member of the TGFβ-family. Notably, when analyzing the scRNA-seq data, TGFβ-signaling was enriched in activated WT DCs compared to activated CD64ΔRamp1 DCs (Fig. S5E). To test the capacity of Activin A to promote DC migration, we utilized an ear explant crawl out assay. Indeed, we found that Activin A enhanced DC migration from the skin (Fig. S5F, G), an effect that was blocked by the TGFβ-receptor family inhibitor Vactosertib (Fig. S5H). *In vivo*, pharmacological inhibition of Activin A signaling reduced DC migration in females and partially in males, effectively eliminating the sex difference (Fig. 5C). Consistent with a direct effect on DCs, CRISPR–Cas9-mediated deletion of *Acvr2a* in hematopoietic cells impaired female DC migration in a cell-intrinsic manner (Fig. 5D). To test, how Activin A promotes DC migration, we incubated skin cells with and without Activin A *ex vivo* and measured the expression of chemokine receptors. We observed Activin A mediated induction of CXCR4 but not CCR7 on skin DCs (Fig. 5E, Fig. S5I), as well as on splenic DCs (unpubl. obs.). Increased CXCR4 levels were functional in enhancing DC migration towards CXCL12 in a transwell assay (Fig. 5F).

We observed CXCL12-GFP expression, the ligand of CXCR4, in lymphatics in activated skin (Fig. S5J) in accord with prior findings^46,47^, supporting the model that CXCR4 can promote the migration of DCs from the skin to the lymph node. Inhibiting CXCR4 pharmacologically reduced DC migration in males and to a greater extent in females, effectively eliminating the sex difference (Fig. 5G). To show that CXCR4-promoted DC migration depended on monocytes, we depleted cells via Gr1 antibody and pharmacologically inhibited CXCR4. As shown before, Gr1 mediated monocyte depletion reduced DC migration, but in CXCR4 blocked mice, Gr1 mediated depletion did not show a significant reduction compared to isotype control (Fig. 5H). Further supporting our model, we observed a marked reduction of CXCR4 expression on activated skin DCs in TrpV1^DTA^ mice compared to WT mice (Fig. 5I). Together, these findings define a neuro–immune axis where CGRP signaling in recruited monocytes induces Activin A production, which in turn primarily promotes DC migration via CXCR4 induction and drives enhanced adaptive immunity in females.

### Androgen signaling in monocytes silences the circuit in males

We next sought to determine what underlies the sex-specific activation of this pathway. Sex differences can arise from either sex chromosome–encoded genetic programs or the influence of sex hormones. To distinguish between these contributions, we first analyzed DC migration using the four core genotypes (FCG) mouse model, which uncouples sex chromosome complement from gonadal sex^48^. The difference in DC migration tracked with gonadal sex rather than sex chromosome complement, with mice bearing ovaries exhibiting greater DC migration than mice bearing testes irrespective of their XX or XY chromosome complement (Fig. 6A). These findings pointed to gonadal rather than sex chromosome–encoded factors as the primary determinant of sexually dimorphic DC migration.

**Fig. 6.**
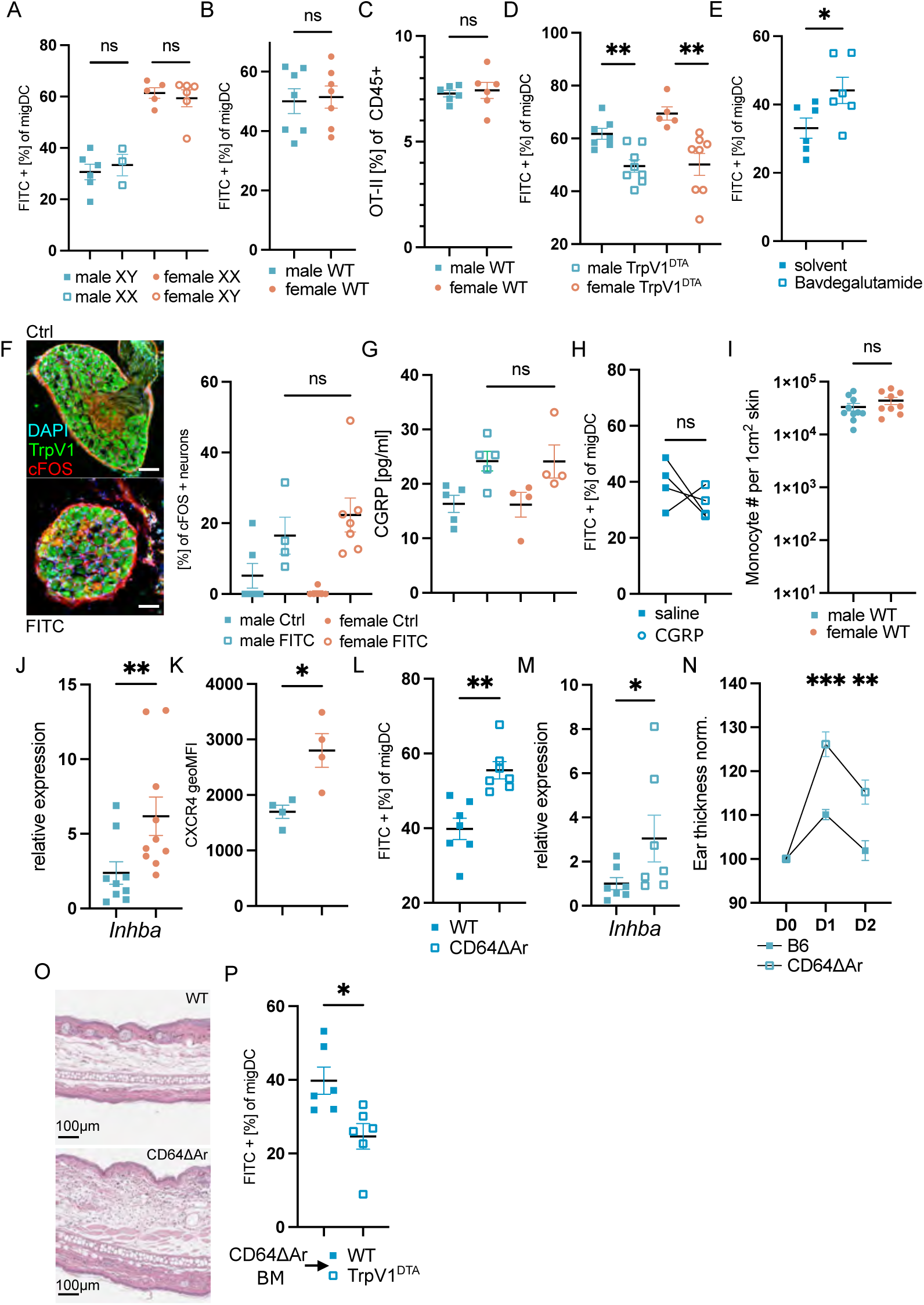
Androgen signaling in monocytes silences the circuit in males. A) Quantification of FITC+ migratory DC in the sdLN 18 h post FITC treatment of four core genotype mice. B) Quantification of FITC+ migratory DCs from the sdLN 18 h post FITC treatment in 3-4 week old WT mice. C) Quantification of OT-II accumulation based on frequency of CD45+ cells at D5 post alum and ova treatment in 3-4 week old WT mice. D) Quantification of FITC+ migratory DCs from the sdLN 18 h post FITC treatment in 3-4 week old WT and TrpV1^DTA^ mice. E) Quantification of FITC+ migratory DC in the sdLN 18 h post FITC treatment in solvent control or Bavdegalutamide treated male mice. F) Representative IF image of DRG 90 min post control (Ctrl) or FITC treatment. TrpV1 (green), cFos (red) and DAPI (blue). Right panel shows quantification of cFos+ DRG neurons 90 min post FITC treatment. G) Quantification of CGRP secretion via ELISA from skin explants after 30 min post control or FITC treatment. H) Quantification of FITC+ migratory DCs from the sdLN 18 h post FITC treatment in male mice with subcutaneous saline or CGRP administration. I) Quantification of Ly6C^high^ monocyte infiltration into the skin 8 h post FITC application. J) Expression analysis of *Inhba* in skin derived Ly6C^high^ monocytes 8 h post FITC application via qPCR. K) Quantification of CXCR4 geoMFI on skin DCs 8 h post FITC application. L) Quantification of FITC+ migratory DCs from the sdLN 18 h post FITC treatment in WT and CD64ΔAr male mice. M) Expression analysis of *Inhba* in skin derived Ly6C^high^ male monocytes 8 h post FITC application. N) Ear thickness measurement at D0, D1, and D2 post challenge phase in male WT and CD64ΔAr mice, normalized to D0 thickness. 8-10 mice per group are displayed. O) Representative H&E histology of ear section collected at D1 post challenge (right). Scale bar: 100 µm. P) Quantification of FITC+ migratory DC in the sdLN 18 h post FITC treatment of BM chimeric mice with CD64ΔAr donor transferred into irradiated male WT or TrpV1^DTA^ hosts. All data were pooled from two to three independent experiments or displays one representative experiment of N= 2-5 mice / group and represented as mean ± SEM. *** *P*<0.001, ** *P*<0.01, * *P*<0.05, ns = non-significant, two-way ANOVA or t-test.

Additionally, we observed in prepubescent mice (3–4 weeks old), the differences in T cell proliferation and DC migration were no longer detectable between males and females (Fig. 6B, C), supporting that sex hormones are required for the emergence of the adult phenotype. These data led us to examine how hormonal signals influence neuronal control of DC migration. In contrast to adult animals, ablation of nociceptor neurons in prepubescent mice reduced DC migration in both sexes (Fig. 6D). Thus, neuronal regulation of DC migration is not intrinsically sex-specific but becomes sexually dimorphic following hormonal maturation. To test this hypothesis, we inhibited androgen receptor (Ar) signaling by treating male mice systemically with Ar Protac degrader for 6 h and analyzed DC migration^49^. Indeed, we observed an increase in DC migration during Ar inhibition, confirming that male sex hormones have an inhibitory effect on DC migration (Fig. 6E). Together, these findings indicate that male sex hormones gate the CGRP-dependent neuronal–monocyte axis, suppressing DC migration and adaptive immunity in males while permitting enhanced responses in females.

We next asked at which level of the discovered axis the sex-specific regulation occurs. To determine whether neuronal activation differs between males and females in our model, we assessed DRG neuron activation following skin FITC challenge. Staining for the activation marker cFOS revealed comparable levels of neuronal activation in both sexes (Fig. 6F). Consistently, CGRP release from skin explants was similar in males and females after treatment (Fig. 6G), indicating that upstream neuronal activation and neuropeptide availability are not limiting factors. Exogenous administration of CGRP failed to enhance DC migration in male mice *in vivo* (Fig. 6H) in contrast to its capability of enhancing DC migration in female TrpV1^DTA^ mice (Fig. 3D), suggesting that male immune cells are refractory to CGRP-mediated signaling.

We therefore examined whether downstream responsiveness to CGRP differs between sexes. Monocyte infiltration, as well as CGRP-receptor transcript expression did not differ between males and females (Fig. 6I, Fig. S6A). However, *Inhba* expression in skin infiltrated monocytes was significantly reduced in males compared to females (Fig. 6J). Analyzing CXCR4 levels on activated skin DCs also revealed a significant difference between males and females (Fig. 6K), although Activin A was able to induce CXCR4 expression similarly on male and female dermal DCs *ex vivo* (Fig. S6B, Fig. 5E). CCR7 expression did not differ between sexes at this early timepoint on activated DCs (Fig. S6C). Together our data indicate that male sex-hormones restrict recruited monocyte ability to promote DC migration.

To test our hypothesis that male sex-hormones modulate monocyte function, we generated mice lacking androgen receptor in monocytes (CD64ΔAr). Analyzing these mice, we could observe an increase in DC migration in males (Fig. 6L), accompanied by an increased expression of *Inhba* in monocytes (Fig. 6M). Furthermore, the CHS response was more severe in these mice compared to WT males (Fig. 6N, O). To further test that the hormone-monocyte axis modulates the neuronal control of DC migration, we generated BM chimeras, by transferring CD64ΔAr BM cells into either WT or TrpV1^DTA^ males. We observed that males lacking the androgen receptor in monocytes showed reduced DC migration in nociceptor neuron deficient mice (Fig. 6P). Together, our data highlight that androgen signaling in monocytes suppresses the neuronal control of DC migration in males by restricting Activin A induction.

In conclusion, our data reveal the existence of a neuroimmune axis, where skin inflammation induces CGRP production by nociceptor neurons, which in turn drives Activin A production by monocytes, upregulating CXCR4 on dermal DCs and thereby promoting their migration to the draining lymph node. In males, this axis is gated by suppressive androgen signaling in monocytes.

## Discussion

Our study identifies an endocrine-neuro–immune circuit that determines the magnitude of cutaneous immunity in a sex-dependent manner. Whereas previous models of sexually dimorphic immunity have largely emphasized direct effects of sex hormones on immune cell numbers and activity^18,50^, our findings reveal an additional layer of regulation in which systemic hormones determine how innate immune cells interpret neuronal signals. Through this mechanism, monocytes integrate endocrine and sensory inputs to control DC migration and the ensuing adaptive immune response. Previous work proposed differences in DC numbers as a basis for sexually dimorphic skin immunity with differences in cutaneous DC abundance controlled by hormone-dependent regulation of GM-CSF production by ILC2s^18^. Our findings extend this framework by demonstrating that sex-biased adaptive immunity can also arise through regulation of DC migration, independently of differences in DC numbers. Together, these observations suggest that sexually dimorphic immunity is shaped not only by immune cell composition but also by the mechanisms through which immune cells integrate activating signals from their tissue environment.

The triangle of sexual-dimorphism, immunity and neurology has been primarily investigated in the context of pain sensation. Tregs^51^, monocytes^35^ and macrophages^34^ have been shown to regulate pain sensation – that is, nociceptor activity – in a sexually dimorphic manner. Here we extend the cross-communication of these systems to how it impacts immune responses. We establish monocytes as the primary integrator of hormonal and neuronal signals that impact the immune response by modulating DC migration. Mechanistically, we show that androgen receptor signaling suppresses CGRP-induced Activin A production, thereby limiting CXCR4-dependent DC migration. Previous *in vitro* studies demonstrated that TGF-β1 can induce CXCR4 expression in human monocyte-derived DCs^52^. Our findings expand this concept by identifying Activin A as an *in viv*o regulator of CXCR4-dependent DC migration and establishing its role in sexually dimorphic immunity. The mechanism of Activin A-induced DC migration we describe differs from that proposed in earlier work^53^ perhaps due to that having been an *in vitro* study. While our findings provide a mechanistic explanation for how Activin A promotes DC migration in females, we do not exclude the possibility that Activin A influences additional aspects of DC biology and migration, as well as androgen signaling impacting monocytes and other immune cells beyond Activin A expression^13,14,50^.

Notably, this circuit was engaged across chemically induced inflammation, microbial challenge and adjuvant-driven immunization. The convergence of these distinct inflammatory contexts on a common neuro–immune pathway suggests that the mechanism described here reflects a general physiological module rather than a stimulus-specific response. More broadly, our findings reinforce the importance of incorporating sex as a biological variable in studies of immunology and, especially neuro-immune regulation.

Several questions remain for future investigation. Although we identify androgen receptor signaling in monocytes as a suppressive mechanism in males, the transcriptional pathways linking androgen signaling to repression of *Inhba* remain unresolved. Also, whether analogous circuits operate in other barrier tissues remains an important question. Neuronal control of immunity and sexually dimorphic immune responses are prominent features of the lung and intestine^13,54,55^, raising the possibility that hormonal gating of neuronal control represents a more general organizing principle of tissue immunity. If so, the implications of this mechanism may extend beyond cutaneous immunity to diverse physiological and pathological contexts in which neuro-immune interactions contribute to host defense and disease.

In this regard, the pathway identified here is notable because multiple nodes—including CGRP, activin-family, CXCR4, and androgen receptor signaling—are pharmacologically tractable^56–59^. These observations raise the possibility that neuronal pathways could be leveraged to modulate adaptive immunity in a sex-aware manner. Nevertheless, neuro-immune interaction is highly context dependent, as shown by the pro-and anti-inflammatory effects of sensory neurons in different contexts^3,11,42,60–63^ and clinical approaches will therefore need to be carefully considered.

Together, our findings identify a monocyte-centered neuro–immune–endocrine circuit that controls DC migration and adaptive immunity in the skin. More broadly, they suggest that sex hormones shape immune responses not only by directly regulating immune cell numbers or activation but also by determining how immune cells interpret information from the nervous system. These findings provide a framework for understanding how interactions among endocrine, neuronal and immune networks generate sex-specific immunity and highlight cross-system communication as a fundamental determinant of tissue immune function.

## Declaration of generative AI and AI-assisted technologies in the manuscript preparation process

During the preparation of this work, the authors used ChatGPT and Claude for text editing and correction purposes. The authors reviewed and edited the output and take full responsibility for the content of all the text.

## Methods

### Animals

All mice were maintained under specific pathogen-free conditions at the University of California, San Francisco. Animals were housed on a 12 h light/12 h dark cycle with ad libitum access to food (PicoLab Mouse Diet 20) and water. Male and female mice aged 7–18 weeks were used as indicated. Littermate and/or co-housed age-matched controls were used for all experiments. C57BL/6J (WT) mice and CD45.1 congenic WT mice (B6.SJL-Ptprca^Pepc^b/BoyJ), OT-II TCR transgenic mice (C57BL/6-Tg(TcraTcrb)1100Mjb/J)^64^, and KikGR transgenic mice (Tg(CAG-KikGR)33Hadj/J)^65^ on a B6 background were bred in-house. Nociceptor neuron-deficient mice (TrpV1^DTA^) were generated by crossing TrpV1-Cre mice (B6.129-Trpv1tm1(cre)Bbm/J; provided by Allan Basbaum, University of California in San Francisco) to Rosa26_lsl-DTA mice (B6.129P2-Gt(ROSA)26Sortm1(DTA)Lky/J). Ramp1^-/-^ mice (B6.129S2(Cg)-Ramp1tm1.1Tsuj/WkinJ)^66^ were purchased from Jackson Laboratories. Conditional deletion of Ramp1 was achieved using Ramp1 fl/fl mice (C57BL/ 6N-Ramp1<tm1c(EUCOMM)Wtsi>/H, provided by Pierangelo Geppetti, University of Florence, Italy) crossed to CD11c-Cre (MGI: 3763248, provided C.A. Lowell University of California, San Francisco), or CD64-Cre (provided by Russell Vance, University of California, Berkeley) or Ms4a3-Cre (C57BL/6J-Ms4a3em2(cre)Fgnx/J)^67^, as indicated. Conditional deletion of the androgen receptor was performed by crossing Ar fl/fl mice (B6.129S1-Artm2.1Reb/J)^68^ to CD64-Cre mice. Conditional deletion of Gnas was performed by crossing Gnas fl/fl mice (Gnastm5.1Lsw/J)^69^ to CD64-Cre mice. CCR2-CreERT2 (Ccr2em1(cre/ERT2)Peng)^70^ mice were crossed to lsl.tdTomato (Gt(ROSA)26Sortm14(CAG-tdTomato)Hze)^71^ and used for histological analysis. CXCL12-GFP (B6.Cg-Cxcl12tm2Tng, provided by T. Nagasawa, Kyoto University)^72^ mice were used for histology. Four core genotype mice (B6.Cg-Tg(Sry)2Ei Srydl1Rlb/JtuJ, provided by Jose Angel Nicolas Avila, University of California in San Francisco)^48^ were used to delineate sex chromosomes and gonadal sex contributions. Ccr2^⁻/⁻^ mice (provided by Q. Tang, University of California, San Francisco) were used in bone marrow chimera experiments to restrict monocyte recruitment, as described below. Cas9-GFP mice were used to generate BM transduction chimeras. Genetically modified strains were maintained on a C57BL/6J background. Prepubescent mice (3–4 weeks old) were used to assess sex hormone-independent effects. All animal experiments were approved by the Institutional Animal Care and Use Committee (IACUC) of the University of California, San Francisco and performed in accordance with institutional guidelines.

### Contact hypersensitivity (CHS)

For sensitization, mice were shaved on the abdomen and treated with 100 µl of 0.5% FITC dissolved in dibutylphthalate:acetone (1:1) on days −6 and −5. On day 0, mice were anesthetized with isoflurane and baseline ear thickness was measured using a digital caliper (Mitutoyo). Mice were then challenged with 25 µl of 0.5% FITC or vehicle control applied to the ear. Ear thickness was measured at 24 h and 48 h post-challenge. For cellular analysis, ears were harvested at 48 h post-challenge and processed for flow cytometry.

### Skin Histology

Ears were harvested 24 h after challenge, fixed in 10% formalin for 24 h, and processed for hematoxylin and eosin (H&E) staining by HistoWiz. For immunofluorescence, skin was shaved, harvested and fixed in 2.5% formalin for 24 h, cryoprotected in 30% sucrose in PBS for 24 h and embedded in OCT and stored at −80°C. Skin tissue was sectioned in 100 μm using a CM3050S cryostat (Leica) and mounted onto Superfrost Plus slides. After rehydration with PBS for 10 min at RT, sections were permeabilized and blocked with PBS containing 1% FCS (Sigma-Aldrich), 1% GCWFS (Sigma Aldrich), 0.3% Triton-X 100 (Carl Roth) and 1% normal mouse serum (Invitrogen) for 60 min at RT. Staining was performed in blocking buffer at 4°C overnight. Sections were washed with PBS and mounted with Aqua-mount. Slides were imaged on a Leica SP8 confocal microscope (Leica) and analyzed using Imaris 9.3.1 (Oxfort Instruments).

## Cell isolation

### Skin

A shaved skin patch of 1 cm^2^ was dissected, minced finely with scissors, re-suspended in 1 ml of digestion media (composed of 0.25 mg/ml Liberase TM, 0.5 mg/ml hyaluronidase and 0.1 mg/ml DNase in RPMI with 1 mM Hepes, and 2% fetal bovine serum), and incubated in a heated shaker at 37°C at 1000 rpm for 60 minutes. Alternatively, 1 ear was separated into ventral and distal halves, minced and digested in the same manner. An additional 1ml of RPMI/Hepes/FBS media containing 1mM EDTA was added and the suspension, vortexed and then strained into a 50 ml conical tube through a 70 μm strainer. Another 10 ml of media was added to thoroughly wash the cell strainer and the suspension was centrifuged, re-suspended in staining buffer (PBS containing 2% FBS) for cell counting and staining. Cell counts were derived from Cytek aurora analyzer and calculated based on acquired volume.

### Lymph node and spleen

Mice were euthanized, spleen and LNs were isolated, spleen was minced into small pieces, and tissues were placed in 1ml digestion buffer (composed of 10 μg/ml collagenase IV and 0.1 mg/ml DNase in RPMI with 1 mM Hepes, and 2% fetal bovine serum). The tissues were rotated at 37°C for 60 minutes. Afterwards, a single cell suspension was generated using a microtube pellet pestle (Fisher Scientific) and the cells were transferred into staining buffer (PBS containing 2% FBS) for staining. Before staining, spleen red blood cells were lysed in RBC lysis buffer.

### Treatment

Mice were shaved with a clipper at the dorsal back skin before treatment. Olcegepant (100 μM, MedChem Express), Qx314 (600μM, Fisher), alpha-CGRP (αCGRP, 250 ng, MedChem Express), Vactosertib (250 μM, MedChem Express), AMD3100 (1 mM, Fisher), and QWF (500 μM, Fisher) were injected subcutaneously in 50 μl at the stated concentrations dissolved in saline or saline with 1% DMSO, or saline with 1% DMSO and 20% PEG 300. FITC was dissolved in a 1:1 mix of dibuthylphtalate:acetone and freshly prepared before treatment and applied topically. αCGRP and Olcegepant were injected 2 h after FITC application, while the other reagents were injected at the time of topical treatment. Bavdegalutamide (50 μg, MedChem), was injected intraperitoneally in saline with 5% DMSO.

### Immunization

Papain (50 μg), Heat Killed *Listeria monocytogenes* (HKLM, 1×10^10^ CFU) generated by incubating at 90°C for 60 min, aluminum hydroxide (Alhydrogel, Invivogen) in a 1:1 mix with saline were injected subcutaneously in a volume of 50 μl with 20 μg ovalbumin added. 25 μl of an 8 mg/ml FITC solution in a 1:1 mix of dibutylphthalate:acetone was applied topically to the skin, in addition to 50 μl subcutaneous injection of saline containing 20 μg ovalbumin.

### Tamoxifen treatments

CCR2-CreERT2 mice were crossed to Ai14 reporters for histological analysis. Tamoxifen (Sigma) was resuspended overnight at 20 mg/ml in corn oil, shaking at 37°C in the dark, and 100 μl was injected intraperitoneally. 1 day later, mice were treated with dibutylphthalate:aceton on the shaved skin and tissue with harvested for histological analysis 24 h later.

### Photoconversion

Mice were shaved and depilated the day before the experiment. To photo-convert the back skin of KikGR-transgenic mice, the mice were anesthetized with isoflurane. Using a Silver LED 415 (Prizmatix), 1cm^2^ of skin was exposed to 415nm violet light for 5 min. Appearance of photo-converted cells in the lymph node and conversion rate of cells in the skin was assessed 18 h post exposure.

### OT-II proliferation assay

Spleens from OT-II-transgenic mice were harvested, a single-cell suspension was created and red-blood cells were lysed. Splenocytes were stained with cell trace violet (CTV, Fisher) dye based on manufactures instructions. 2×10^6^ CTV-labeled splenocytes were transferred i.v. into recipient mice. 18 h later mice were immunized subcutaneously with 20 μg Ova and either 25 μl topical application of 8 mg/ml FITC in dibutylphthalate:Acetone, 50 μl alum, or 1×10^10^ CFU HKLM in 50 μl saline. FITC and HKLM treated mice were analyzed 2 days post immunization for CTV dilution. Alum treated mice were analyzed 5 days post immunization and OT-II cells were quantified and restimulated to analyze their cytokine profile.

### Cytokine measurement

sdLN from immunized mice were isolated and a single-cell suspension was generated. 100.000 cells were plated in a 96-well plate and stimulated using PMA-ionomycin. Golgistop (BD Biosciences/Fisher) was added to the incubation media. Cells were incubated for 5 h at 37°C. Afterwards cells were washed using FACS buffer and surface markers were stained. Golgistop was added to the antibody mix. Then, cells were fixed and lysed using CytoFix buffer (BD). Cytokines were stained over night at 4°C in Perm/Wash buffer.

### Ear crawl assay

Mice were euthanized and ears were harvested. Ventral and dorsal side were separated and placed floating on 1.5 ml migration media (RPMI containing 0.5% fatty acid free BSA, 1 mM HEPES). The tissue was incubated for 48 h at 37°C. Afterwards, migrated cells were collected from the media and analyzed and quantified via flow cytometry. Activin A (100 ng/ml) and Vactosertib (25 μM) were added to the migration media where indicated.

### Transwell migration assay

Spleens were harvested and splenocytes were isolated by digesting the tissue for 1 h in 1 ml digestion buffer (composed of 10 μg/ml collagenase IV and 0.1 mg/ml DNase in RPMI with 0.5% fatty-acid free BSA, 10 mM Hepes). Red blood cells were lysed using RBC buffer and cells were counted and resuspended in migration medium (0.5% fatty-acid free BSA, 10 mM Hepes in RPMI) at 2×10^7^ cells/ml. Cells were incubated for 4 h at 37°C with or without Activin A (100 ng/ml, Thermo Fisher). Afterwards cells were washed and used for Transwell migration assay without the presence of Activin A. Transwell filters (6 mm insert, 5 μm pore size, Corning) were placed on top of each well, and 100 μl containing 2×10^6^ cells were added to the transwell insert. No ligand (Nil), or CXCL12 (50 ng/ml, Peprotech) were added to the media at the bottom of the transwell. The cells were allowed to migrate for 3 h at 37°C with 5% CO_2_ and the cells in the bottom well were stained and counted by flow cytometry.

### BM chimera

Mice, where indicated, were lethally irradiated with 900 rads gamma-irradiation (split dose separated by 4 h) and then i.v. injected with 5×10^6^ BM cells. To generate mixed BM-chimera, BM cells from each donor mouse were counted using a hemocytometer and 5×10^6^ BM-cells were transferred in total with a mixing ratio of 50:50. Mice were analyzed 6-8 weeks post reconstitution.

### Antibody depletion

To deplete cell types, antibodies were injected into the peritoneal cavity 1d before treatment of the mice. Gr1 (50 μg, BioXcell) and Ly6G (100 μg, BioXcell) were injected intraperitoneally. Depletion efficacy was tested in the skin or spleen, respectively.

### Hot plate assay

Mice were brought into the testing room 30–60 minutes before the procedure for acclimation. The surface of the hot plate was cleaned and heated to a maximum temperature of 53°C. One mouse at a time was placed on the hot plate and the latency to show a nocifensive response such as hind paw withdrawal, licking, stamping, leaning, or jumping is recorded. The mouse was immediately removed once this response was observed. If there was no response within about 40 seconds, the test was terminated, and the mouse removed from the hotplate to prevent heat-related injury.

### ELISA

Amount of secreted CGRP was measured using ELISA assay (LSBio). 0.5 cm^2^ skin was harvested 30 min post treatment and placed on 1 ml RPMI (containing HEPES and Pen/Strep). The tissues were incubated for 4 h at 37°C. The supernatant was collected and processed based on manufactures instructions.

### DRG isolation and histology

Mice had their entire back skin depilated 72 h before the experiment to allow sensory acclimation. 300 μl FITC was applied to the back skin and 90 minutes later, mice were perfused with PBS and then 4% paraformaldehyde. DRGs were dissected and fixed in 4% paraformaldehyde at 4°C for 1 hour, cryoprotected in 30% sucrose in PBS for 24 hours and embedded in OCT and stored at −80°C. DRG tissue were cut in 30 μm at a cryostat (Leica) and mounted onto Superfrost Plus slides. Primary antibodies used were rabbit anti-c-Fos and guinea pig anti-TRPV1. Secondary antibodies used were Alexa488-conjugated donkey anti-guinea pig IgG, Alexa647-conjugated goat anti-rabbit IgG. Slides were mounted with Fluoroshield™ with DAPI, imaged on a Leica SP8 confocal microscope (Leica) and analyzed using Imaris 9.3.1 (Oxfort Instruments).

### sgRNA mediated deletion

For CRISPR-Cas9 mediated gene deletion, single-guide RNA (sgRNA) sequences were cloned into the pTR-MSCV-IRES-Thy1.1 or pTR-MSCV-IRES-BFP vector. SgRNA sequences were selected using Benchling’s CRISPR Guide tool and 2 guides were designed for Acvr2a. As control empty vectors were used. To produce retroviruses, Platinum-E packing cells (Cell Biolabs) were grown to 75% confluency in a 10 cm tissue culture dish and transfected with 10 μg plasmid and 25 μl of Lipofectamine 2000 (Thermo Fisher) in OptiMEM following manufacturer’s protocol. ViralBoost (Alstem) was added to the media 16 h later and retrovirus containing supernatants were collected 48 and 72 h after transfection, filtered through a 0.45 μm filter to remove debris, and concentrated 10 fold with Retro-X concentrator (Takara). For BM transductions, constitutive CAS9-GFP expressing donor mice were injected i.p. with 3 mg of 5-fluorouracil (Sigma), BM was collected at day 5 and cultured with complete DMEM supplemented with 20 ng/ml interleukin-3 (IL-3), 50 ng/ml IL-6, and 100 ng/ml stem cell factor. BM cells were spin-infected (1000 g, 2 h, 32°C) at day 1 and day 2 and transferred into irradiated recipients after the second transduction.

sgRNA-sequences

Acvr2a sg1:

GAGAAAAGAGGCACCAGTGG

Acvr2a sg2:

TGGGCAGGTTGGTACCCGGG

### scRNA-sequencing

Mice were treated with FITC on the dorsal flank as described above. 8 h post treatment, skin cells were isolated and stained as described above. The samples were sorted using a BD FACSAriaII with a 100 μm nozzle into RPMI medium supplemented with 10% FBS. CD64+, CD45+ cells were sorted and dermal DCs based on CD11c and MHC2 expression were sorted separately. After sorting, cells were counted and CD64+ cells were mixed with cDC in a 9:1 ratio. The 4 samples were processed using 10x on-chip multiplexing (OCM) chip based on manufacturers instructions by the UCSF Institute for Human Genetics Sequencing Core.

Transcripts captured in all the cells encapsulated with a bead were uniquely barcoded using a combination of a 16 bp 10x Barcode and a 10 bp unique molecular identifier (UMI). cDNA libraries were generated using the Chromium™ Single Cell 3’ Library & Gel Bead Kit v3 (10x Genomics) following the detailed protocol provided by the manufacturer. Libraries were sequenced with the NovaSeq 6000 platform (S1 Cartridge, Illumina) in 150 bp paired-end mode. The library was aligned and demultiplexed using CellRanger software (version 7.0.0)^73^. Aligned reads were used to quantify the expression level of mouse genes and generation of gene-barcode matrix. Subsequent data analysis was performed using Seurat R package (4.5.2)^74^. Quality control was performed, and viable cells were selected by excluding cells with features lower than 800 and above 7000, as well as cells having more than 7% of mitochondrial transcripts and more than 70000 transcripts. 2000 most variable genes were used for downstream analysis to calculate principal components, after log-normalization and scaling. Principle component analysis (PCA) was used for dimensionality reduction and to visualize a uniform manifold approximation and projection (UMAP) of the identified clusters. P-values comparing gene expression of clusters and samples were calculated with the FindMarker function in Seurat. Gene list was generated using an adjusted p-value cutoff <0.05 and expression in the indicated cluster cutoff >25%. Differential gene expression comparing WT_FITC and CD64ΔRamp1_FITC samples of cluster Monocyte_1 was calculated using DEseq2 statistical test. Pathway analysis was performed using EnrichR^75–77^ with standard settings.

### Flow cytometry

The cells were stained in FACS buffer (PBS containing 2% FBS) with FC block (anti-CD16/32, BioXcell) to block nonspecific binding, antibody mix, and with Zombie NIR (Biolegend) for 30 min at 4°C to exclude dead cells. The following antibodies and clones were purchased from Biolegend: CD45.2 BV605, AL700 (104), CD45.1 AL700 (A20), CD24 PB (M1/69), Ly6C BV650 (HK1.4), Epcam BV711 (G8.8), CD86 PerCP-Cy5.5 (GL-1), CD11c PE, AL647, SparkNIR6 (N418), Sirpa PE-Dazzle594 (P84), CD64 PE-Cy7 (W18349C), TCR Va2 FITC (B20.1), CD301b APC (URA-1), CD103 PE-Cy5 (2E7), XCR1 AL647 (ZET), Ly6G APC-Cy7 (1A8). The following antibodies and clones were purchased from BD: CD45.2 BUV395 (104), CD4 BUV615 (GK1.5), TCRβ BUV737 (H57-597), CD19 BUV563 (1D3), IA/IE BUV805 (M5/114.15.2), CD11b BUV496 (M1/70), CD44 BUV496 (IM7). The cells were run on Aurora (Cytek) or Symphony (BD) and data were analyzed using FlowJo software (BD).

### Chemokine Receptor staining

The cells were stained in FACS buffer (PBS containing 2% FBS) with Fc block (anti-CD16/32, BioXcell) to block nonspecific binding, and CCR7 Biotin (4B12, BioLegend), CXCR4 BV421 (2B11, BD) for 45 min at 37°C. Aftwards cells were stained with SA-PE (ThermoFisher) and additional surface antibodies for 30 min at 4°C.

### Analysis of public RNA-seq data

Publicly available RNA-seq data were retrieved from GEO (GSE247477 and GSE250456) and *Inhba* expression was retrieved from the dataset as provided online. *Acvr1b* and *Acvr2a* expression data was retrieved from immgen.org.

### Quantitative RT-PCR

For assessment of gene expression total RNA was extracted using RNeasy kit and reverse-transcribed using M-MLV reverse transcriptase. qPCR was performed using Power SYBR Green with an Applied Biosystems StepOnePlus instrument. Data were analyzed with the comparative Ct (2−ΔCt) method, using *Hprt* as housekeeping gene control.

Primer sequences:

Murine Inhba fw AACCACTACCGCATGAGGGG

Murine Inhba rev TTTCTCTGGGACCTGGCGAC

Murine Calcrl fw CCTGGGACGGATGGCTATG

Murine Calcrl rev ACTTTCTCATGCGTGCTGTTAT

Murine Ramp1 fw CCTCTGCTTACCTCTGAGATTG

Murine Ramp1 rev ATCTGTGCAGTCTTCCTTGGAGT

Murine Hprt fw AGGTTGCAAGCTTGCTGGT

Murine Hprt rev TGAAGTACTCATTATAGTCAAGGGCA

## Statistical analysis

All experiments were performed 2 or more independent times, with the exception of the scRNA-seq experiment. Statistical analyses were performed using GraphPad Prism software. Unpaired or paired two-tailed Student’s t test was used to compare two groups, for multiple group comparison two-way ANOVA was used. P values smaller than 0.05 were considered statistically significant. Error bars represent mean ± standard error of the mean (SEM). scRNA-seq experiment was analyzed using Seurat package in R. Investigators were not blinded to group assignment during experimental procedures or analysis.

## Materials availability

Requests for resources and reagents should be directed to and will be fulfilled by J.G.C.

## Data availability

scRNA-sequencing data is deposited at GEO.

## Acknowledgement

We thank Pierangelo Geppetti for Ramp1 floxed mice, Russell Vance and Bernard Malissen for CD64-Cre mice. TrpV1-Cre mice were provided by Allan Basabaum. We thank current and former Cyster lab members for helpful discussion and feedback. We especially thank Zachary Earley for providing HKLM, the UCSF Parnassus Flow Core CoLabs for their assistance in sorting, the UCSF Parnassus Functional Genomics Core CoLabs for their assistance in generating scRNA-seq data. K.K. was supported by DFG Walter Benjamin postdoctoral fellowship and is currently supported by Cancer Research Institute Irvington Postdoctoral Fellowship. J.G.C. is an Investigator of the HHMI. This work was supported in part by NIH Grant AI 40098.

## Competing interests

J.G.C. is a consultant for Lycia Therapeutics and Dren Bio.

## Author contributions

K.K. and J.G.C. designed research; K.K., T.L.B.-S., J.A., and Y.X. performed research; K.K. and J.G.C. analyzed data; and K.K. and J.G.C. wrote the manuscript with input from all authors.

## Supplementary Figure legends

**Fig. S1.**
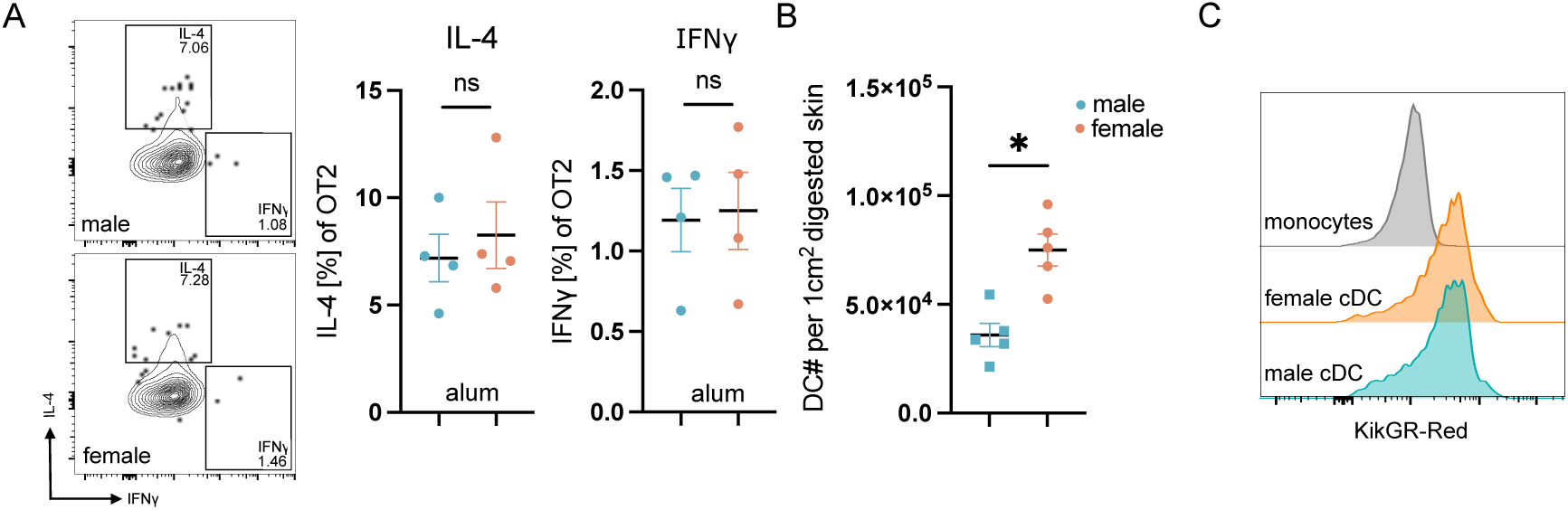
Characterization of female-biased cutaneous immune responses, Related to. Fig. 1 A) Representative flow cytometry plot of transferred OT-II cell isolated from the sdLN at D5 post immunization with alum and ova (left), quantification of cytokine expression of transferred OT-II cell isolated from the sdLN at D5 post immunization with aluim and ova (middle, right). B) Quantification of absolute DC numbers per 1cm^2^ of digested skin from WT mice. C) Representative flow cytometry plot of KikGR-Red photoconverted cells of either female monocytes and DCs, as well as male DCs, isolated from the skin 18 h post photoconversion and HKLM immunization. The majority of monocytes are recruited after the 5 min photoconversion period and are used as an unconverted control. All data were pooled from two to three independent experiments or displays one representative experiment of N= 3-4 mice / group, and represented as mean ± SEM. * *P*<0.05, ns = non-significant, t-test.

**Fig. S2.**
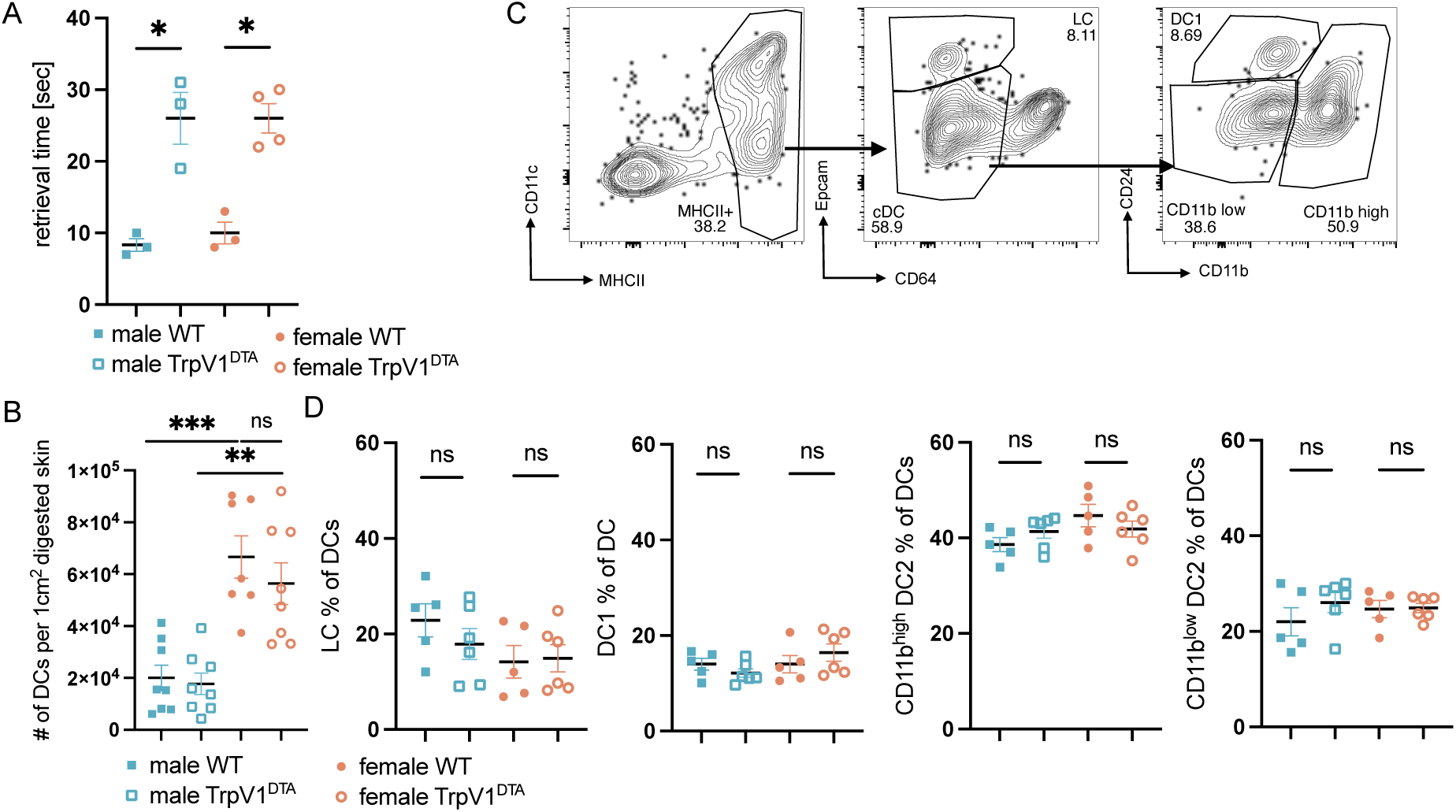
Nociceptor ablation does not alter skin dendritic cell abundance or composition, Related to. Fig. 2 A) Quantification of retrieval time of WT and TrpV1^DTA^ mice in a hot plate assay. B) Quantification of absolute DC numbers per 1cm^2^ digested skin of WT and TrpV1^DTA^ mice. C) Representative flow cytometry plot displaying gating strategy to identify skin DC subsets. LC = Langerhans cells; cDC = conventional DCs; DC1 = type 1 conventional DCs; CD11b low = CD11b low expressing type 2 conventional DCs; CD11b high = CD11b high expressing type 2 conventional DCs. D) Quantification of skin DC subsets. All data were pooled from two to three independent experiments or displays one representative experiment of N= 3-4 mice / group, and represented as mean ± SEM. *** *P*<0.001, ** *P*<0.01, * *P*<0.05, ns = non-significant, two-way ANOVA or t-test.

**Fig. S3.**
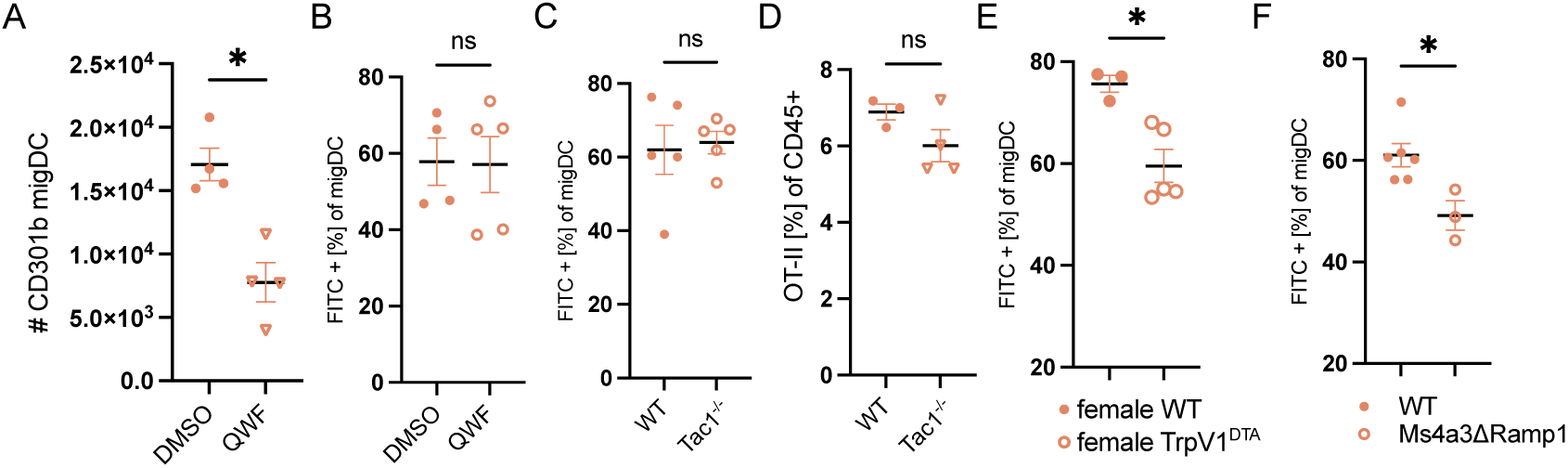
Hapten-mediated dendritic cell migration is independent of Substance P, Related to. Fig. 3 A) Quantification of CD301b+ migratory DC in the sdLN 18 h post papain immunization in female mice. B) Quantification of FITC+ migratory DC in the sdLN 18 h post FITC treatment in female mice. C) Quantification of FITC+ migratory DC in the sdLN 18 h post FITC treatment in WT and Tac1^-/-^ female mice. D) Quantification of OT-II accumulation based on frequency of CD45+ cells at D5 post alum-and ova treatment in WT and Tac1^-/-^ female mice. E) Quantification of FITC+ migratory DC in the sdLN 18 h post FITC treatment of BM chimeric mice with WT donor transferred into irradiated female WT or TrpV1^DTA^ hosts. F) Quantification of FITC+ migratory DC in the sdLN 18 h post FITC treatment in WT and Ms4a3ΔRamp1 female mice. All data were pooled from two to three independent experiments or displays one representative experiment of N= 2-4 mice / group, and represented as mean ± SEM. * *P*<0.05, ns = non-significant, t-test.

**Fig. S4.**
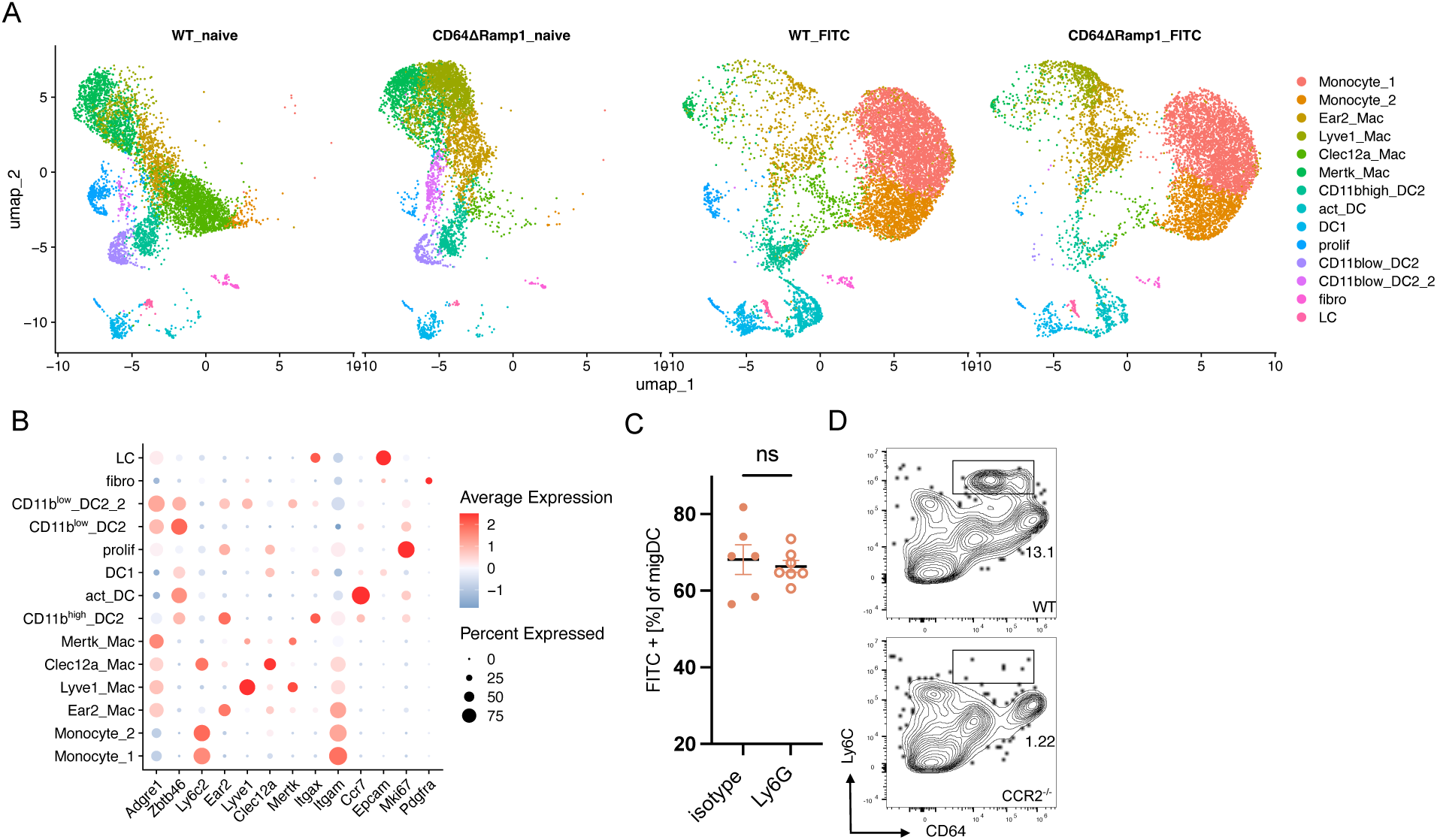
Recruited monocytes mediate CGRP-dependent dendritic cell migration, Related to. Fig. 4 A) UMAP projection displaying the clustering of sorted CD64+ immune cells based on scRNA-seq data, color represents cluster identity, split by sample. B) DotPlot displaying expression level of cluster defining genes. C) Quantification of FITC+ migratory DC in the sdLN 18 h post FITC treatment in anti-Ly6G treated female mice. D) Represented flow cytometry plot displaying infiltration of Ly6C^high^ monocytes into the skin 8 h post FITC application in WT and CCR2-deficient female mice. All data were pooled from two to three independent experiments or displays one representative experiment of N= 3 mice / group, and represented as mean ± SEM. ns = non-significant, t-test.

**Fig. S5.**
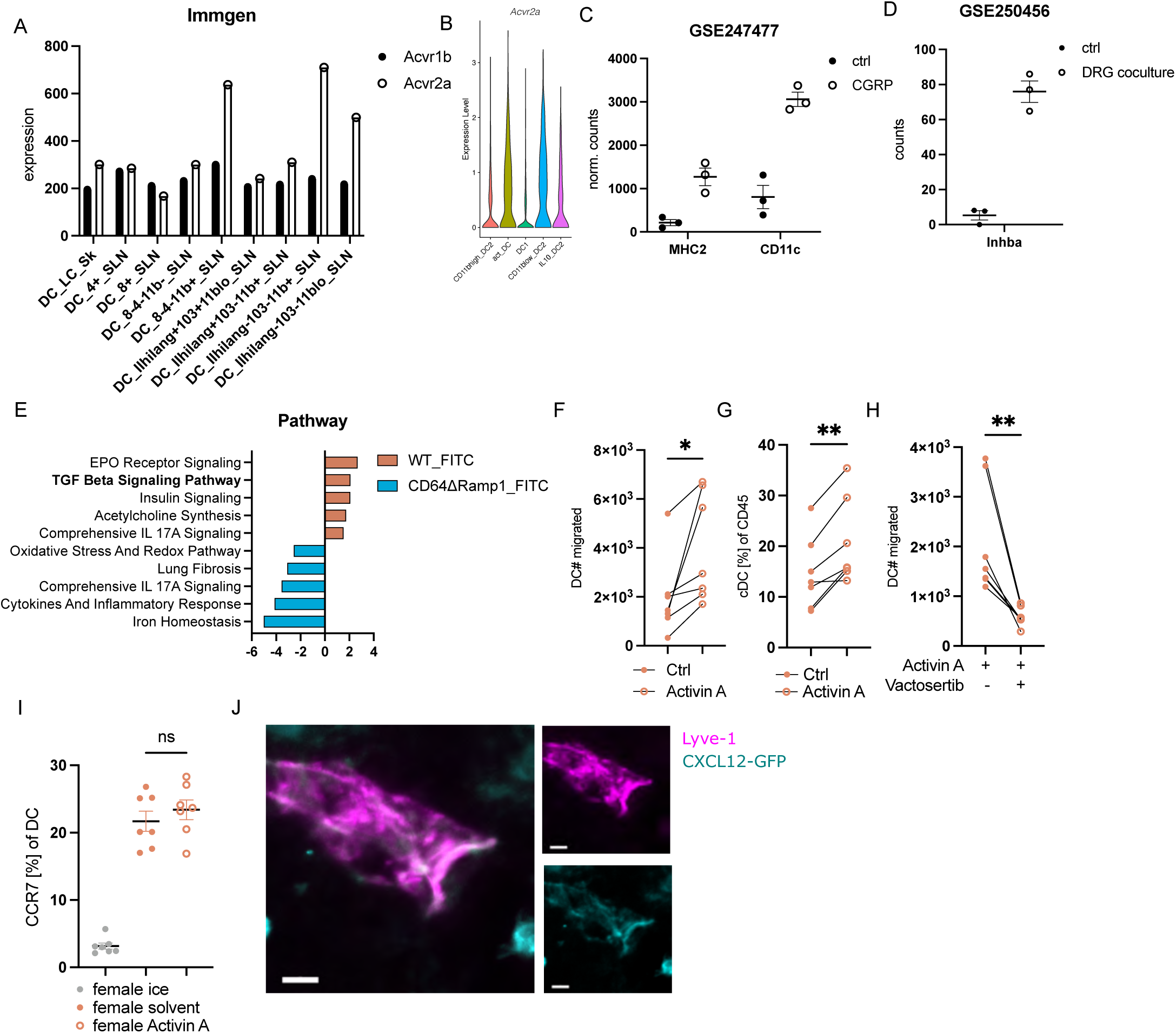
Activin A promotes CXCR4-dependent dendritic cell migration, Related to. Fig. 5 A) Expression level of *Acvr1b* and *Acvr2a* in sdLN DC. Data derived from Immgen. B) Quantification of *Acvr2a* expression across DC clusters from scRNA-seq data. C) Normalized counts level of *Inhba* expression in MHC2+ and CD11c+ macrophages. Data derived from GSE247477. D) Counts level of *Inhba* expression in BMDM cells with or without DRG coculture. Data derived from GSE247477. E) Pathway analysis of DE genes comparing WT and CD64ΔRamp1 cells from act_DC cluster. Top 5 pathways for each sample are displayed. F) Quantification of migrated DC numbers from ear-crawl-out assay after 48 h comparing solvent control and Activin A treated female samples. G) Quantification of migrated DC frequency from ear-crawl-out assay after 48 h comparing solvent control and Activin A treated female samples. H) Quantification of migrated DCs from ear-crawl-out assay after 48 h comparing Activin A treated samples with Activin A and Vactosertib treated female samples. I) Quantification of CCR7 expression on female skin DCs after *ex vivo* treatment with Activin A for 4 h. J) Representative IF image if CXCL12-GFP expression in female skin 24 h post dibutylphthalate application. CXCL12 (cyan) and Lyve-1 (magenta). Scale bar: 10 µm. All data were pooled from two to three independent experiments or displays one representative experiment of N= 3-4 mice / group, and represented as mean ± SEM. ** *P*<0.01, * *P*<0.05, ns = non-significant, t-test.

**Fig. S6.**
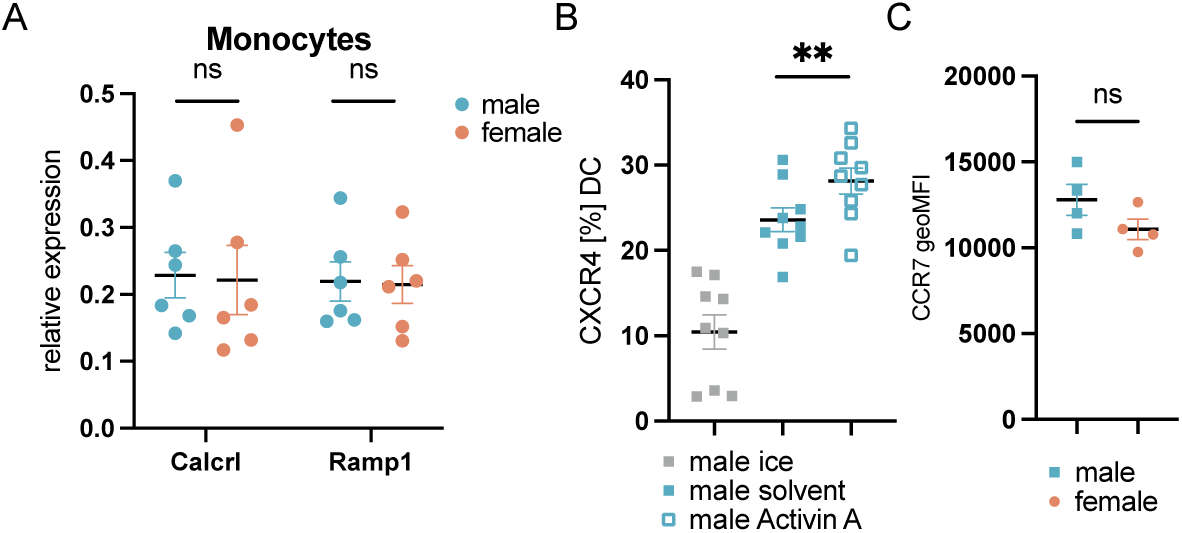
Sex differences in dendritic cell migration arise upstream of dendritic cells, Related to. Fig. 6 A) qPCR analysis of *Calcrl* and *Ramp1* expression in splenic monocytes. B) Quantification of CXCR4 expression on male skin DCs after *ex vivo* treatment with Activin A for 4 h. C) Quantification of CCR7 geoMFI in skin DCs 8 h post FITC application. All data were pooled from two to three independent experiments or displays one representative experiment of N= 3-4 mice / group, and represented as mean ± SEM. * P<0.05, ns = non-significant, t-test.

